# Regeneration of Plants from DNA-free Edited Grapevine Protoplasts

**DOI:** 10.1101/2021.07.16.452503

**Authors:** Simone Scintilla, Umberto Salvagnin, Lisa Giacomelli, Tieme Zeilmaker, Mickael A. Malnoy, Jeroen Rouppe van der Voort, Claudio Moser

## Abstract

CRISPR-Cas technology has widely extended the application fields of genome editing in plant breeding, making possible specific and minimal mutations within a genetic pool. With respect to standard genome editing technologies, CRISPR-Cas machinery can be introduced in the form of ribonucleoproteins (RNPs), thus avoiding the introduction of exogenous DNA into cells. The interest on the application of DNA-free delivery into plant cells is constantly increasing, especially in the case of valuable woody plants elite varieties where CRISPR-Cas9 technology would preserve their genotype, while still resulting into targeted genetic modifications. The use of single cells fits well the requirements of New Breeding Technologies, by ensuring both the CRISPR-Cas DNA-free delivery as RNPs and, since every plant will be regenerated from a single edited cell, the absence of chimerism. However, the use of protoplasts cell culture from woody plants is generally hampered by low editing efficiencies and an unsuccessful regenerative process.

We here describe a successful DNA-free methodology to obtain fully edited grapevine plants, regenerated from protoplasts obtained from V. vinifera cv. Crimson seedless L. embryogenic callus. The transfected protoplasts were edited on the Downy Mildew susceptibility gene VvDMR6-2. The regenerated edited plants exhibited homozygous deletions of 1bp or 2bp, and homozygous insertion of 1bp.

## Introduction

The genome editing technology allows to modify cellular DNA with a high level of precision. In particular, with the advent of CRISPR-Cas9 (*Clustered Regularly Interspaced Short Palindromic Repeats-*Cas9) technology, the application fields of genome editing have been widely extended. This system is based on the recognition of the DNA-editing site through a complementary RNA sequence and the Cas nuclease–mediated DNA double-strand break, which makes the insertion, deletion or even just the modification of one nucleotide possible. Therefore, especially in the case of genetic improvement of woody plants (e.g. grapevine or apple) elite varieties, the CRISPR-Cas9 technology ensures their genotype preservation, while resulting in targeted genetic modifications. The CRISPR-Cas components can be introduced inside a cell either in the form of nucleic acids (i.e. DNA/mRNA coding for the entire system), or in the form of ribonucleoprotein (RNP) complex. While DNA can integrate into the genome and mRNA is affected by its intrinsic instability, direct cellular delivery of RNPs opens up to attractive scenarios as it potentially embodies a robust methodology leading to a specific and minimal mutation with no trace of exogenous DNA (Woo et al., 2015). With this perspective, the interest on the application of a CRISPR-Cas technology to plants, potentially of better acceptance by consumers as compared to classic GMOs, is constantly increasing (Saleh et al., 2021).

Three main strategies have been proposed so far to deliver the CRISPR-Cas system into plant cells. 1) The use of engineered *Agrobacterium tumefaciens* which easily overcomes the plant cell wall. However, this strategy employs exogenous plasmid DNA containing portions of DNA from Agrobacterium, which gets integrated in the cellular DNA upon transformation. In the case of woody plants, exogenous DNA can only be removed by outcrossing, resulting in changes to the genetic background. An alternative approach to segregation, successfully applied to many crops including woody plants, consists in the molecular excision of the T-DNA (Dalla Costa et al., 2020), by which the exogenous DNA is almost entirely removed. However, the minimal residual foreign DNA remaining is likely incompatible with the current strict GMO regulations of many countries. 2) Particle bombardment uses nanoparticle bullets loaded with biological material to shoot plant tissues, thus overpassing the cell wall barrier, and releasing the nanoparticles-loaded biological cargo to induce genome editing. This strategy fits well with DNA-free strategy requirements. Nevertheless, various physical parameters severely affect the efficiency of this approach. In particular, as not all the cells would be hit by the bullets, the downstream regeneration process may give rise to chimeric plants. 3) An alternative solution is the temporary removal of the cell wall for an effective delivery of biological material into single cells. According to this strategy, the cell wall is digested enzymatically, thus providing a single, “naked” plant cell (i.e. protoplast) delimited by the plasma membrane. Under favorable conditions, cellular delivery of RNPs can be easily achieved by means of PEG infiltration, electroporation or lipofection. After 2-3 days the cell wall is restored, and further cellular divisions will lead to the regeneration of a whole plant. As every plant will be regenerated from a single, edited cell, this methodology guarantees genetic homogeneity of the final product.

## Results and discussion

The single-cell strategy is the most promising among the three proposed ones, and it has been successfully tested on many crops. However its application to woody plants such as grapevine and apple is hampered by low editing efficiencies and an unsuccessful regenerative process, which usually stops after few cellular divisions. For grapevine, the scientific literature reports successful regeneration of whole plants from protoplasts obtained from embryogenic callus (Reustle et al., 1995). Hence, upon development of genome editing techniques, embryogenic callus constitutes the best candidate tissue from which obtaining edited and potentially regenerating protoplasts (Bertini et al., 2019; Malnoy et al., 2016; Tricoli, 2019). Among these works, two reported the regeneration of whole plants from wild-type protoplasts (Bertini et al., 2019; Tricoli, 2019), but to date, none reported regeneration of whole plants from edited protoplasts.

The present work provides a new methodology to i) obtain protoplasts from grapevine embryogenic callus; ii) edit the protoplasts at a target gene through CRISPR-Cas9 and iii) regenerate fully edited grapevine plants. The methodology was successfully applied to *V. vinifera* cv. Crimson seedless *L*., a popular table grapevine variety from which highly regenerative callus is easily obtained.

Protoplasts were isolated from 1 g of Crimson s. grapevine embryogenic callus in 12 ml of enzymatic mixture composed by 1% (w/v) cellulase Onozuka R-10, 0.2% (w/v) hemicellulase Sigma, 0.3 % (w/v) macerozyme R-10 dissolved in Gamborg B5 (including vitamins and 0.45 M mannitol) under sterile conditions. Next, the suspension was poured into a 90 mm Petri-dish and left on a tilt-shaker at 25°C for 16 hours in darkness. After incubation, the suspension was filtered through a 60 μm nylon sieve, and protoplasts were collected by centrifugation at 80 g for 4 min, without brake. Next, the collected protoplasts were washed twice in an osmotically adjusted MMG medium (Malnoy et al., 2016).

To avoid contaminations of intact single cells after the digestion phase, protoplasts were further purified layering 2 ml of the protoplasts suspension on 8 ml of a 16 % w/v sucrose aqueous solution. Healthy protoplasts were collected at the interphase upon centrifugation (90 g, 4 min, no brake) (**Fig. 1A**). Protoplasts were successfully transfected with the RNP complex composed by Cas9 protein (Thermofisher) and a sgRNA (guide: GGAGGATTGGAGGGCCACTC, **Fig. 1B**) to edit the *VvDMR6-2* gene by PEG infiltration, as described elsewhere (Malnoy et al., 2016). The *DMR6* (*Downy Mildew Resistant* 6) gene was initially described in *A. thaliana* as susceptibility gene to different pathogens (Zeilmaker et al., 2015). Loss-of-function mutations in the grapevine DMR6 orthologs *VvDMR6-1* and *-2* and, especially *VvDMR6-2*, were significantly more represented in Downy Mildew-resistant genotypes (Pirrello et al., 2021), thus making *VvDMR6-2* a good candidate for a CRISPR-Cas9 knock-out aiming to obtain downy mildew tolerant grapevine plants.

**Figure 1.**
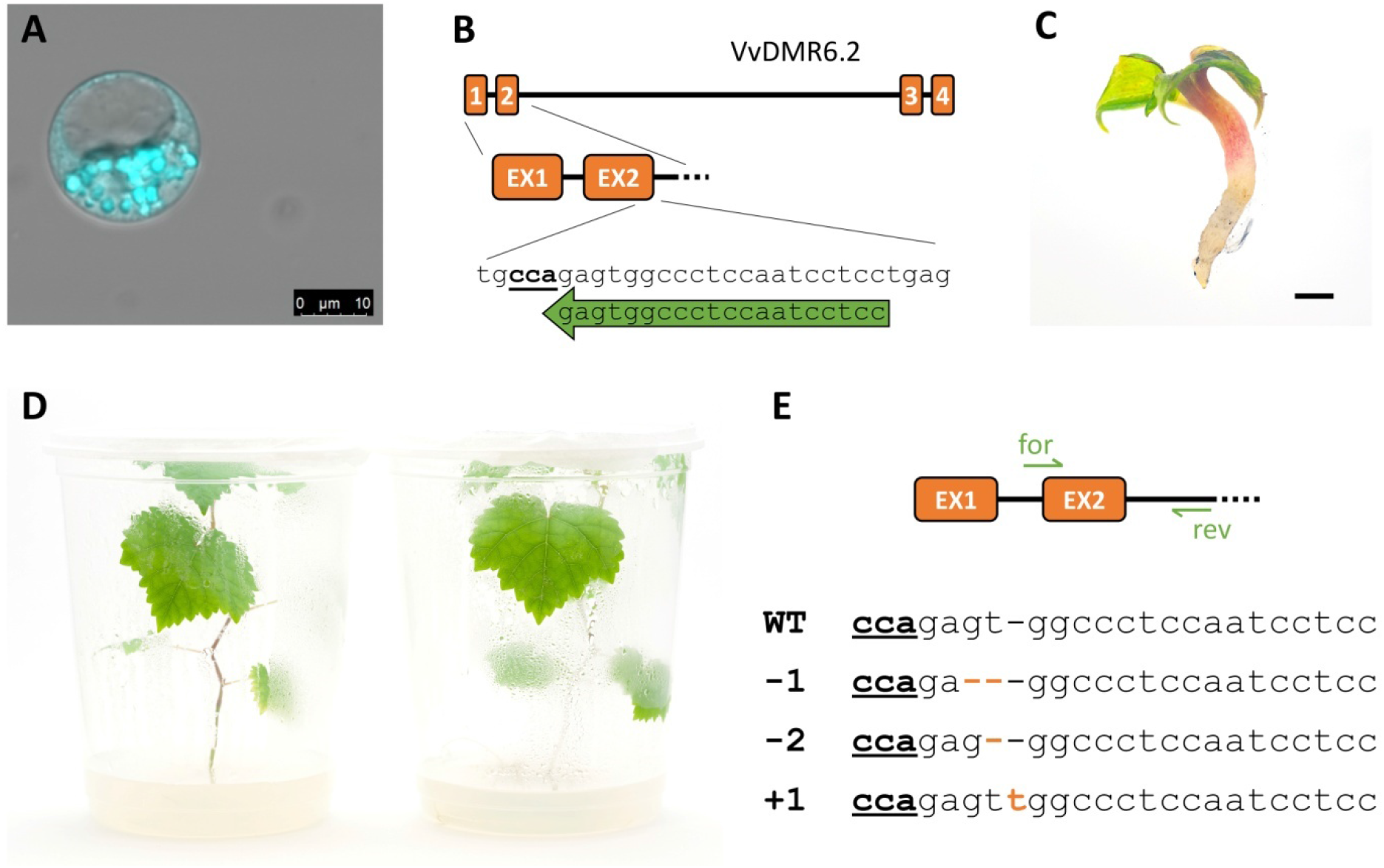
DNA-free editing and regeneration of grapevine plants from protoplasts. (A) FDA-staining of a viable protoplast from the embryogenic callus digestion mixture. (B) Gene structure (exons and introns) of *VvDMR6-2* and sgRNA target site. (C) Embryo obtained after 4 months of alginate disk culture (2 months in liquid NN culture and 2 months in solid GS1CA, respectively). Black bar corresponds to 2mm. (D) WT (left) and edited (right) plants obtained from protoplast cultures after 5-6 months. (E) Editing profiles as confirmed by Sanger sequencing on the amplicons. WT sequence (1^st^ row) is compared with indel mutants (−1bp, -2bp and +1bp, 2^nd^, 3^rd^ and 4^th^ row, respectively).

Alginate disks containing either wild-type or edited protoplasts were cultured and stimulated for the formation of mini-calli in a 50 mm Petri-dish filled with 5 ml of a NN-based cultivation medium (Nitsch and Nitsch, 1969) optimized for regeneration (Nitsch and Nitsch medium including vitamins, 88 mM sucrose, 300 mM glucose, 1g/l charcoal, 0.93 μM kinetin, 2.22 μM 6-BAP and 10.7 μM NAA). Cultures were stored at 24 °C in permanent darkness and the liquid culture medium was changed weekly. After 2 weeks, the glucose concentration was progressively diminished by 25% per week, and after 4 further weeks no glucose was present in the regenerative culture medium. The disks were transferred on solid GS1CA culture medium, enriched with 300 μM glutathione. After 3 to 4 weeks upon transfer to solid medium, the growth of the first embryos was observed. These embryos were then transferred on NN solid medium and stored at 16/8 light/dark photoperiod (80-100 μmol m^-2^ s^-1^), at 24 °C. Successful plant development was obtained within 3 to 4 months from the start of the whole procedure (**Fig. 1C, D**).

*VvDMR6-2* gene editing on plants regenerated from transfected protoplast was confirmed by Sanger sequencing on amplicons obtained from genomic DNA (DMR6for, “TCACTGGTCACATCCACACC”, DMR6rev “AAAATGATGCGGGAGGACAT”, 435bp).Seven plants showed homozygous editing out of the eight plants tested. In particular, the observed mutations were the homozygous deletion of 1bp or 2bp, and a homozygous insertion of 1bp (**Fig. 1E**).

The methodology proposed is potentially applicable to any grapevine - either table or wine – variety and for the editing of any target gene, thus representing a rapid and accurate process for the genetic improvement of any trait for which the genetic bases are known.

## Acknowledgements

This work was supported by Fondazione Caritro, Bando Ricerca e Sviluppo 2018, funding grant for the project TRADING (Transfer of molecular probes into plant cells for genome editing and transcriptional profiling, 2018.0261), 2018/2020.

## Author coontributions

S.S., C.M., M.A.M. and J.R.V. designed the research. S.S. designed the experiments. S.S. and U.S. performed the experiments and collected the samples. L.G. provided embryogenic callus. L.G. and T.Z. designed the sgRNA. The manuscript was written by S.S. and U.S. and edited by S.S., U.S. L.G., T.Z., M.A.M., J.VR. and C.M. All authors read and approved the manuscript.

## Competing interests

S.S., U.S., L.G. and T.Z. are named inventors on a patent application pertaining to the technology filed by Fondazione E. Mach and Enza Zaden. All other authors have no competing interests.

